# A genetically encoded sensor for real-time monitoring of poly-ADP-ribosylation dynamics in-vitro and in cells

**DOI:** 10.1101/2024.06.11.598597

**Authors:** Alix Thomas, Kapil Upadhyaya, Daniel Bejan, Hayden Adoff, Michael Cohen, Carsten Schultz

## Abstract

ADP-ribosylation, the transfer of ADP-ribose (ADPr) from nicotinamide adenine dinucleotide (NAD^+^) groups to proteins, is a conserved post-translational modification (PTM) that occurs most prominently in response to DNA damage. ADP-ribosylation is a dynamic PTM regulated by writers (PARPs), erasers (ADPr hydrolases), and readers (ADPR binders). PARP1 is the primary DNA damage-response writer responsible for adding a polymer of ADPR to proteins (PARylation). Real-time monitoring of PARP1-mediated PARylation, especially in live cells, is critical for understanding the spatial and temporal regulation of this unique PTM. Here, we describe a genetically encoded FRET probe (pARS) for semi-quantitative monitoring of PARylation dynamics. pARS feature a PAR-binding WWE domain flanked with turquoise and Venus. With a ratiometric readout and excellent signal-to-noise characteristics, we show that pARS can monitor PARP1-dependent PARylation temporally and spatially in real-time. pARS provided unique insights into PARP1-mediated PARylation kinetics in vitro and high-sensitivity detection of PARylation in live cells, even under mild DNA damage. We also show that pARS can be used to determine the potency of PARP inhibitors in vitro and, for the first time, in live cells in response to DNA damage. The robustness and ease of use of pARS make it an important tool for the PARP field.

---

PARP1 is a critical first responder to various types of cell DNA damage^1^. The binding of PARP1 to damaged DNA leads to its activation via long-range allostery. Active PARP1 catalyzes ADP-ribosylation of itself and other protein targets (e.g., histones) using nicotinamide adenine dinucleotide (NAD^+^) as a substrate^2-3^. PARP1-mediated ADP-ribosylation leads to the recruitment of DNA damage response (DDR) proteins and, ultimately, DNA repair^4^. DDR-defective cancer cells are uniquely and profoundly sensitive to the loss of PARP1, referred to as synthetic lethality. This finding inspired the clinical development of PARP1 inhibitors, five of which are FDA-approved for the treatment of DDR ovarian and breast cancer^5^.

For years, it was thought that PARP1 predominately generates polymers of ADP-ribose, a process called poly-ADP-ribosylation (PARylation) on glutamate and aspartate residues of protein targets. Yet recent studies show that PARP1 catalyzes mono-ADP-ribosylation (MARylation) on serine residues of protein targets^6^, a process significantly enhanced by the co-factor protein HPF1^7-8^. The initial site of serine MARylation may become a starting point for further serine PARylation; however, a recent proteomics study suggests that serine in PARP1 targets is predominately MARylated and not PARylated^9^, suggesting that in cells, PARylation occurs predominately on glutamate/aspartate.

Like other PTMs, glutamate/aspartate PARylation and serine MARylation are reversible. ADP-ribose hydrolase 3 (ARH3) is the only known serine MARylase in cells^10-11^, whereas poly-ADP-ribose glycohydrolase (PARG) is the predominant PARylase in cells. The rapid reversal (minutes timescale) of PARylation by PARG under DNA damage conditions is critical for faithful DNA repair; knockdown of PARG or inhibition of PARG activity results in defects in DNA repair, underscoring the critical role of PARylation in the DNA damage response^12^.

The transient nature of PAR in cells makes it challenging to study PARylation using conventional methods such as Western blotting. An effective approach to tracking the spatiotemporal dynamics of PARylation in live cells is using a genetically encoded sensor. A typical sensor design is based on a domain that recognizes PAR with high specificity and selectivity. Such PAR-binding domain-based sensors have been described^13-15^. However, they suffered from a low signal-to-noise ratio and were only shown to detect PAR levels qualitatively under strong PARP1 activation conditions.

Here, we describe the design and characterization of a highly sensitive and specific Forster Resonance Energy Transfer (FRET) based sensor, which we call pARS, that dynamically monitors PARP1-dependent PARylation in vitro and in live cells (Figure 1). We observed PARP1-mediated PARylation kinetics on the seconds time scale, which revealed sigmoidal kinetics suggesting allosteric modulation of PARP1. pARS could semi-quantitatively measure changes in PARylation in live cells in response to increasing DNA damage. Finally, we find that pARS can be used for determining PARP inhibitor potency in live cells, demonstrating its potential for screening PARP inhibitors in a cellular context.

**Figure 1.**
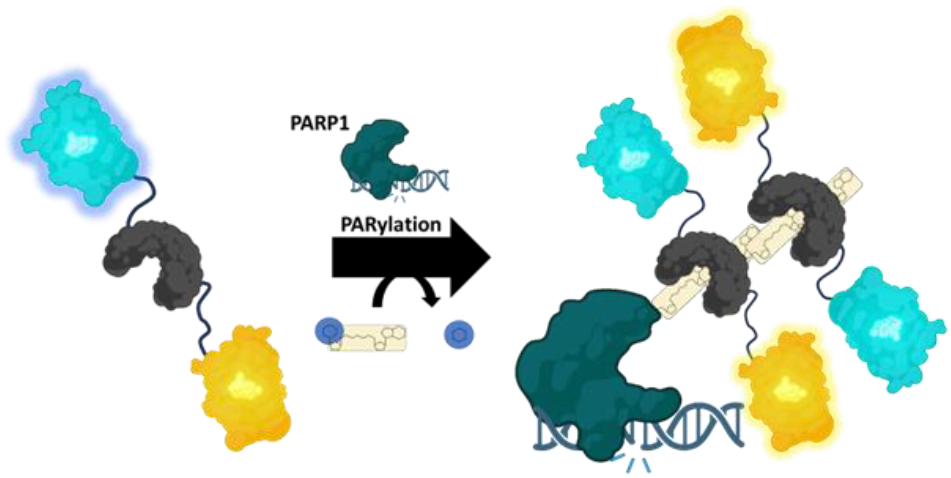
Sensor design of pARS. Generated using BioRender

To detect PARP1-mediated PARylation, we designed a FRET-based sensor that contains the specific PAR-binding domain WWE from the ubiquitin E3 ligase RNF146, sandwiched between a well-established pair of fluorescent proteins with mTurquoise (mTurq) as a donor and the yellow fluorescent protein mVenus as acceptor^16^ (Figure 1)^17-18^. When PARP1 is PARylated in vitro, the sensor oligomerizes on PAR units resulting in a strong increase in FRET compared to baseline level (Figure 1 and 2A). The addition of trypsin leads to the cleavage of the sensor and a loss of the observed FRET. Because the FRET change depends on the oligomerization of the sensor onto PAR chains, the molecular ratio of pARS over PARylated PARP1 is critical in setting the dynamic range. An excess of the sensor can lead to a reduced dynamic range, attributable to residual, unbound sensor. At the same time, an insufficient sensor-to-PAR ratio may cause signal loss due to the increased distance between each sensor molecule. We show here that a molecular ratio of 50/1 pARS-to-PARP1 is optimal to maximize the dynamic range of the sensor (Figure S1). Surprisingly, we noticed that upon adding DNA without PARP1, pARS exhibited an increase in FRET comparable to the addition of auto-PARylated PARP1 (Figure S2A). We investigated if pARS can directly bind to DNA, which would lead to an unwanted increase in FRET. pARS and DNA were incubated at different pARS concentrations in the presence or absence of DNAse. Starting at 5 μM pARS, we observed a shift in DNA migration and smearing of the sensor, which was prevented by adding DNAse, indicating that pARS binds to DNA in the micro-molar range in-vitro (Figure S2B,C). Next, we incubated a recombinant WWE domain with DNA and found that WWE binds to DNA with a similar affinity to pARS. The WWE-Y145A mutant, which cannot bind to PAR, also loses its ability to bind to DNA, suggesting that WWE binds DNA and PAR. Accordingly, we prevented unspecific FRET increase of pARS by using lower DNA concentration when performing PARP1 reactions.

**Figure 2.**
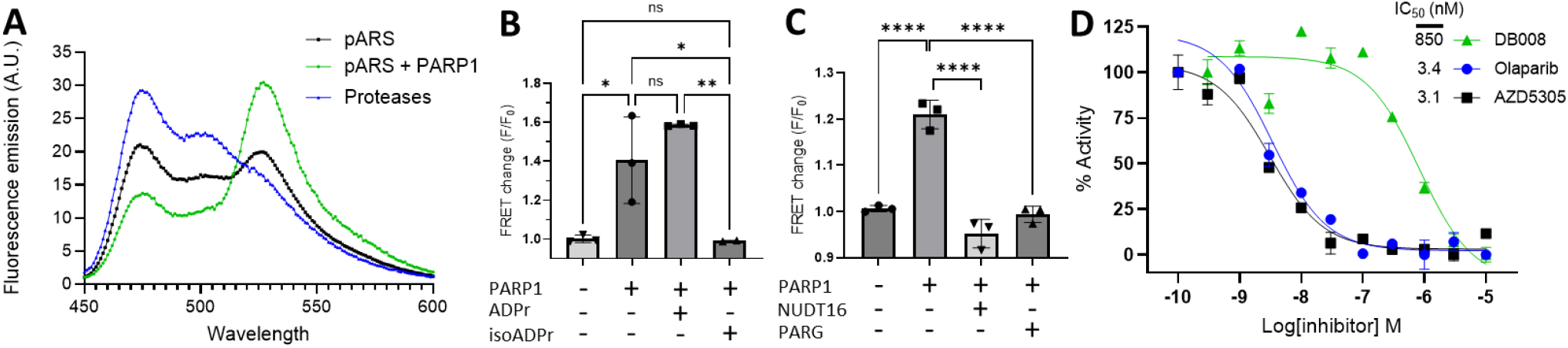
**A**. Fluorescence emission spectrum of pARS (5 μM) excited with 440 nm (+/-10) light with or without PARP1 (100 nM). **B**. FRET change of pARS upon PARP1 activity with or without pre-incubation with 10 μM ADPr or isoADPr. **C**. FRET change of pARS upon PARP1 activity with or without the remover enzymes NUDT16 and PARG. **D**. pARS FRET reporter assay to assess potency of Olaparib, AZD5305, and DB008 against PARP1 (10 nM) in a plate reader. *p < 0.05, **p < 0.01, ***p < 0.005, ****p < 0.001 (One way Anova). At least three independent experiments.

We next assessed the ability of pARS to distinguish MAR from PAR. The smallest known unit of PAR recognized by the WWE domain is isoADPr, which is the unique repeating unit of PAR^17-18^. We synthesized isoADPr by an improved synthetic path^19-20^ (see the supplement) and incubated pARS with isoADPr before adding PARylated PARP1. isoADPr effectively blocked the PARylated PARP1-mediated FRET change. In contrast, pre-incubation with ADPr, which does not bind to the WWE domain^17^, did not significantly impact the PARylated PARP1-mediated FRET change (Figure 2B). These results demonstrate the specificity of pARS for PAR versus MAR. pARS specificity was further tested using enzymes that can degrade PAR: NUDT16, a pyrophosphatase, and PARG, an O-glycohydrolase^21^. Treatment of PARylated PARP1 with these enzymes led to a complete loss of FRET (Figure 2C), further confirming the selectivity of pARS for detecting PAR. Finally, we utilized pARS to determine the potency, in a 384 well plate format, of the clinically approved PARP1 inhibitors Olaparib and AZD5305 as well as DB008^22^, a low potency PARP1 inhibitor. We obtained IC_50_ of 3.4 and 3.1 nM for Olaparib and AZD5305 respectively, and 850 nM for DB008, in alignment with previous reports^23-24^ (Figure 2D).

To accurately monitor PARP1 kinetics in vitro, we added the sensor to PARP1 before initiating the PARP1 reaction. This allowed us to follow PARP1 activity in real time on the second time-scale. We found that increasing NAD+ concentrations increased the rate and magnitude of the FRET change. Interestingly, we observed a delayed increase in the reaction rate and the optimal fitting curve of our data corresponded to an allosteric/sigmoidal model, suggesting allosteric modulation of PARP1 (Figure 3A). As expected, the maximum reaction velocity increased with NAD^+^ concentration following a hyperbolic fit. We determined a K_m_ for PARP1 of 8.62 μM (Figure 3B), consistent with prior published Km values on full length PARP1^25^. Intriguingly, decreasing the temperature during the PARP1 auto-PARylation reaction appears to increase either the total amount of PAR chains or their length, as reflected by the gradual higher maximum product concentration when the reaction proceeded at a lower temperature (Figure S3). Finally, we investigated PAR removal by PARG in real-time (Figure 3C). The addition of PARG rapidly reduced FRET following an exponential decay model. Inhibition of PARG using the selective small molecule PARG inhibitor PDD00017273 (PDD), prevented the PARG-mediated FRET decay. By examining the kinetics of PARP1 and PARG with resolution at the seconds timescale, we hope to facilitate and expand the range of applications compared to previously established PAR reporters^13, 26-27^, and open new avenues for characterizing the regulation of PARP1-mediated PARylation by modulators and inhibitors, and how mutations in PARP1 and PARG can impact their activities.

**Figure 3.**
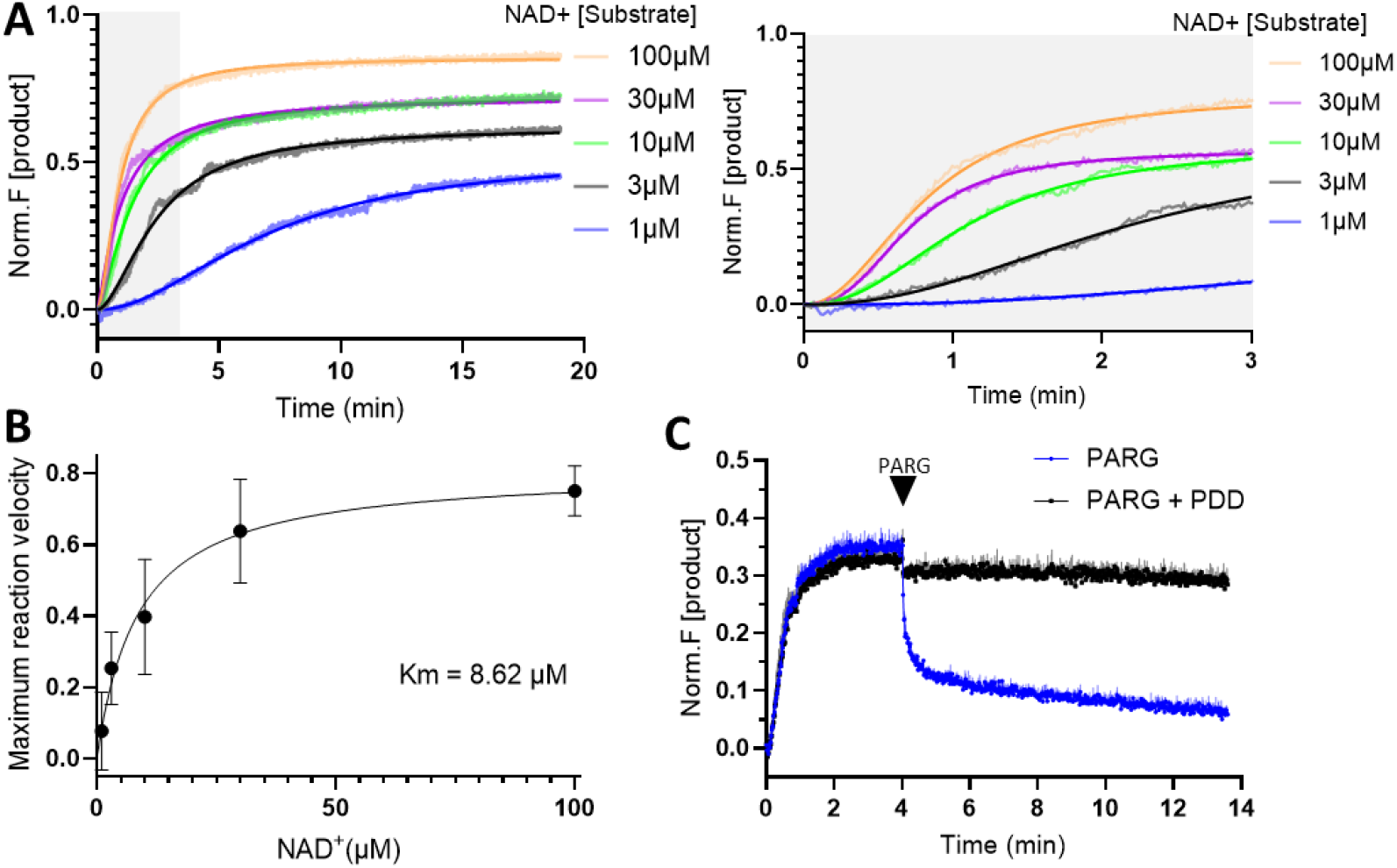
**A**. PARP1 (5 nM) dependent PAR formation under various NAD^+^ concentrations. Curve was fitted using an allosteric/sigmoidal non-linear regression model. In grey is a zoom of figure A showing sigmoidal fit in early time points. **B**. Normalized reaction velocities from **A**. for determining Km via Michaelis-Menten kinetics of PARP1. **C**. PARP1 (5 nM) dependent PAR formation at 10 μM NAD^+^ (37°C) with the addition of PARG (20 nM) in the presence or absence of PDD (1 μM). Data is shown as Mean ± SEM of two independent experiments.

Having established pARS as a robust sensor for monitoring PARP1-dependent PARylation in vitro, we next sought to evaluate pARS in live cells. We expressed pARS in HeLa cells and performed live-cell FRET imaging. Multiple nuclear location sequences (NLS) were added to the sensor on its N- and C-terminus to achieve complete nuclear expression of pARS (Figure 4A). Upon treatment of cells with 1 mM H_2_O_2_, which induces DNA damage, we observed an increase in FRET (7.16 fold increase relative to standard deviation) followed by a gradual decrease back to baseline within 20 min. Treatment with the PARP1/2 inhibitor Olaparib (1 μM) fully abolished the response, while adding the PARG inhibitor PDD (1 μM) potentiated the increase in FRET and prevented the decay of the signal, consistent with the notion that PARG is the major PARylase in cells (Figure 4B). Alternative induction of DNA damage using methyl-methanesulfonate (MMS) led to an increase in FRET almost identical to H_2_O_2_ (Figure 4C, Figure S4A). Next, we determined if PARP1 was the major PAR writer in cells. Beyond PARP1, PARP2 is a closely related family member that also PARylates proteins in response to DNA damage. We expressed pARS in WT, PARP1 KO, and PARP2 KO cells and treated with H_2_O_2_. PARP2 KO cells showed the same H_2_O_2_-induced FRET increase as WT cells; by contrast, the H_2_O_2_-induced FRET increase was abolished entirely in the PARP1 KO cells (Figure S4B). These results show that in U2OS cells, the PARylation response after DNA damage is primarily dependent on PARP1 with minimal contribution of PARP2. This confirmed previous observations of PARP2 mainly catalyzing the synthesis of branched PAR chains^28^. Together, these results demonstrate that our sensor can reliably follow the spatiotemporal dynamics of PARylation mediated by PARP1 in response to DNA damage in live cells. To confirm our in vitro experiments that pARS functions via oligomerization of several sensor molecules, we generated variant “homo” versions of pARS with either two mTurquoise or two mVenus fluorescent proteins on each end of the WWE domain. We observed an increase in FRET after treatment with H_2_O_2_ (Figure 4D), albeit with lower sensitivity, suggesting that the mechanism of action is similar in vitro and in cells.

**Figure 4.**
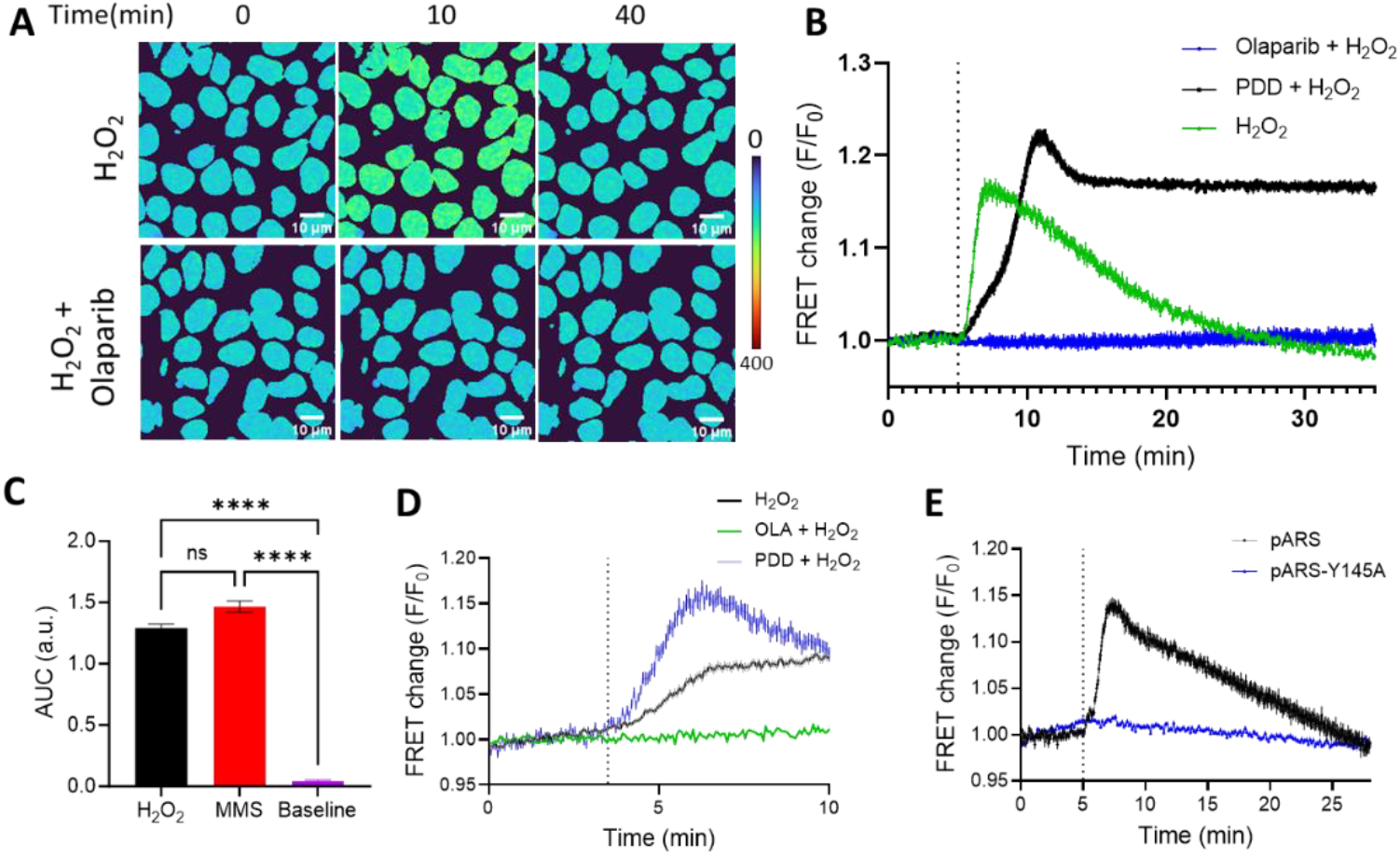
**A**. FRET ratio (CFP/YFP) of HeLa cells transfected with pARS at different time points after treatment with 1 mM H_2_O_2_ with or without PARP1 inhibitor Olaparib. **B**. Average of 27, 24 and 26 HeLa cell traces showing FRET changes of pARS after treatment with 1 mM H_2_O_2_, with or without pretreatment with PDD or Olaparib. **C**. FRET changes of pARS after treatment with 1 mM H_2_O_2_ or 10 mM MMS. **D**. Average of 119, 54 and 32 traces of HeLa cells co-expressing homoTurq-pARS and homoVenus-pARS after treatment with 1 mM H_2_O_2_, with and without pretreatment with Olaparib or PDD. **E**. FRET changes of pARS in 23 cells or the “dead” pARS-Y145A sensor in 72 cells after treatment with 1 mM H_2_O_2_. Data are shown as Mean ± SEM of three (A,B,C) or two (D,E) independent experiments. *p < 0.05, **p < 0.01, ***p < 0.005, ****p < 0.001 (One way Anova)

Additionally, we showed that pARS binds to the site of DNA damage after irradiation with a 375 nm laser similar to the previously reported probes, GFP-WWE and ddGFP-WWE^13, 29^ (Figure S5). Mutating Tyr145 to alanine in the WWE domain of pARS, leading to loss of PAR binding, abolished the FRET changes of the sensor in live cells^17^ (Figure 4E). Taken together, our results show that the FRET increase of pARS upon DNA damage in live cells is driven by the oligomerization of the sensor on PAR.

Treatments such as millimolar concentrations of H_2_O_2_ or MMS induce massive DNA damage within cells but are often the standard used for detecting PARP1-mediated PARylation in cells using more conventional methods such as Western blotting. Given the excellent signal-to-noise of our sensor, we wanted to know if we could detect lower levels of PARylation in live cells using milder DNA damage conditions. We therefore used a 375 nm laser combined with different concentrations of Hoechst, a dye potentiating DNA damage induction by UV irradiation (Figure 5A). By subjecting HEK 293 cells stably expressing pARS to minimal irradiation (0.1% laser power), we characterized the range of PARP1 activity, from saturation with 10 μM Hoechst to a 10% increase of maximum FRET in the presence of 0.2 μM Hoechst (Figure 5B,5C). We did not observe any increase in FRET after laser irradiation in cells transfected with pARS-Y145W, demonstrating the absence of photobleaching potentially contributing to the increase in FRET (Figure S6). These results demonstrate the tunability of the sensor and its ability to semi-quantitatively monitor changes in PARylation dynamics upon minor changes in the DNA damage response in live cells.

**Figure 5.**
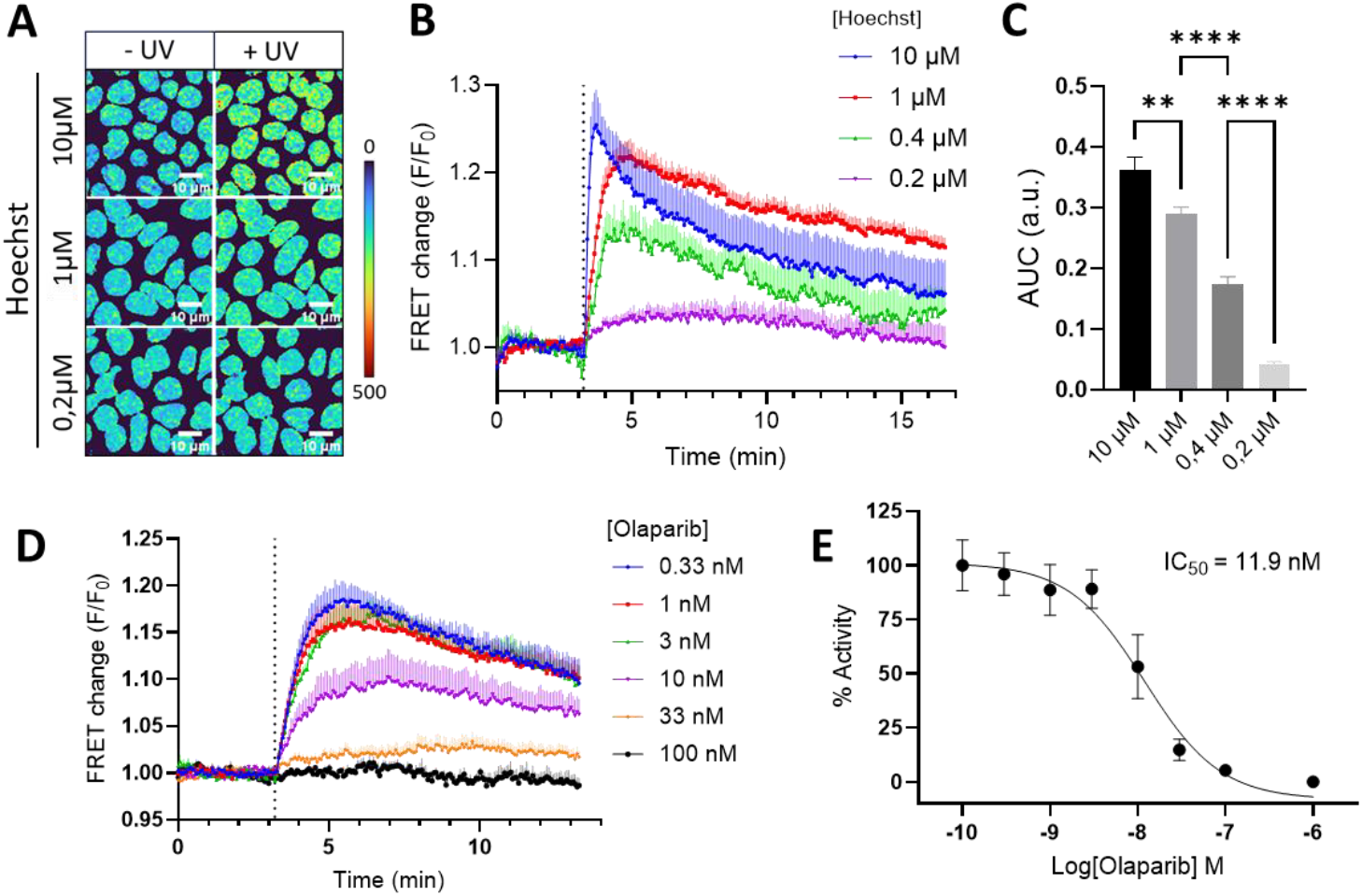
**A**. FRET ratio (CFP/YFP) before and after UV irradiation of HEK 293 cells stably expressing pARS. Cells were treated with 10, 1, or 0.2 μM of Hoechst 5 min before the experiment. **B**. FRET changes of pARS over time as in **A**. after irradiation with 375 nm laser for 1 second. **C**. Bar graph representing the area under the curve (AUG) of each trace in **B**. between 3.2 and 5 min. **D**. FRET changes of pARS stably expressed in HEK 293 pretreated with 1 μM Hoechst and irradiated with a 375 nm laser for 1 sec. Cells were treated with various concentrations of Olaparib. **E**. Dose response curve assessing Olaparib potency in live cells. Values were derived from area under the curve values of Figure 4D traces. Data is shown as Mean ± SEM of three (A,B,C) or two (D,E) independent experiments. *p < 0.05, **p < 0.01, ***p < 0.005, ****p < 0.001 (One way Anova)

Lastly, we sought to demonstrate the utility of pARS for determining the potency of PARP1 inhibitors in live cells. We used 375 nm irradiation in the presence of 1 μM Hoechst, which induced about 80% of the dynamic range of the sensor. This ensured that the FRET signal was not saturated. Incubation of HEK 293 cells stably expressing pARS with increasing concentrations of Olaparib led to a dose-dependent decrease in the FRET response (Figure 5D,5E). We then calculated the area under the curve for each dose-response. We obtained an IC_50_ of Olaparib in live HEK293 cells of 11.9 nM, which fits in the median of literature values ranging from 3nM to 250 nM obtained using Western blotting ^22, 25, 30^. Together, these results demonstrate that pARS is useful for evaluating PARP1 inhibitor potency in live cells.

In summary, by utilizing intermolecular FRET, pARS efficiently detects PARP1-mediated PARylation, enabling semi-quantitative measurements in vitro and live cells. This novel approach allows the characterization of PARP1 auto-PARylation kinetics at unprecedented second-scale resolution, potentially advancing our understanding of PARP1 modulation by cofactors and inhibitors.

By monitoring PARP1 auto-PARylation in real-time in-vitro, we unveiled intriguingly delayed kinetics. We believe this is due to PAR acting as an allosteric activator, consistent with recent studies^31^. Structural studies have demonstrated that PARP1 activity is influenced by the type of DNA breaks, shifting it from cis to trans-autoPARylation^27, 32-34^. This factor could contribute to our observed kinetics.

Leveraging the dynamic range of the pARS sensor, we established a novel live-cell method for quantitative measurement of PARP1-dependent PARylation. This approach enabled precise IC_50_ determination for PARP inhibitors and promises to facilitate the assessment of their dissociation constants (k_off_) in live cells, a critical factor for determining pre-clinical efficacy^35^. This offers significant advancement over previous qualitative probes^13-14^.

The interplay between PARylation and MARylation-mediated by PARP1 is considered pivotal for developing next-generation inhibitors. Novel genetically encoded biosensors capable of quantifying MARylation alongside pARS could revolutionize our understanding of PARP1’s regulatory mechanisms^29^.

## ASSOCIATED CONTENT

### Supporting Information

The authors have cited additional references within the Supporting Information^19-20^.

The Supporting Information is available free of charge on the ACS Publications website

## AUTHOR INFORMATION

## Author Contributions

Alix Thomas designed and performed experiments, analyzed data, interpreted the results, and wrote the original draft of the manuscript. Kapil Upadhyaya designed and performed experiments. Daniel Bejan designed and performed experiments. Hayden Adoff performed experiments. Michael Cohen and Carsten Schultz contributed the experimental design, analyzed the data, and edited the manuscript.

## Notes

The authors declare no competing financial interests.

## ACKNOWLEDGMENT

C.S. acknowledges financial support from OHSU and an endowment donated by Helen Jo and Bill Whitsell. C.S. is a recipient of a Mercator Fellowship from the DFG, connected to Transregio 186. M.S.C. acknowledges funding from the NIH (2R01NS088629).

## TABLE OF CONTENTS

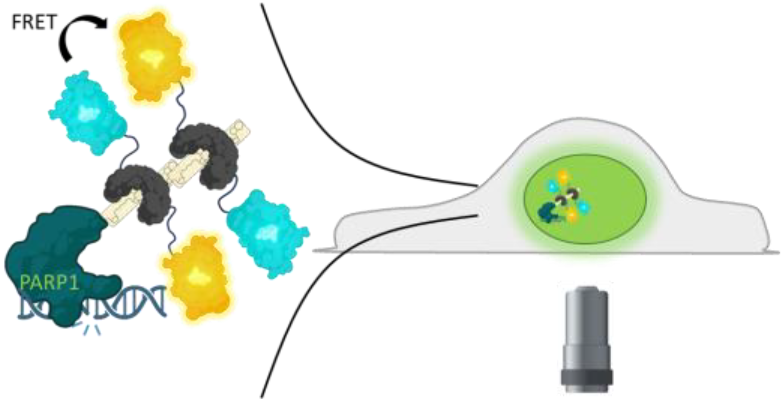

PARP1 is the primary DNA damage-response writer responsible for adding a polymer of ADPR to proteins (PARylation). Real-time monitoring of PARP1-mediated PARylation is critical for understanding the regulation of this unique PTM. Here, we describe a genetically encoded FRET probe (pARS) for semi-quantitative monitoring of PARP1-dependent PARylation temporally and spatially in real-time in vitro and in live cells.

