## Supporting information for "A genetically encoded sensor for real-time monitoring of poly-ADP-ribosylation dynamics in-vitro and in cells"

**SUPPORTING FIGURES**

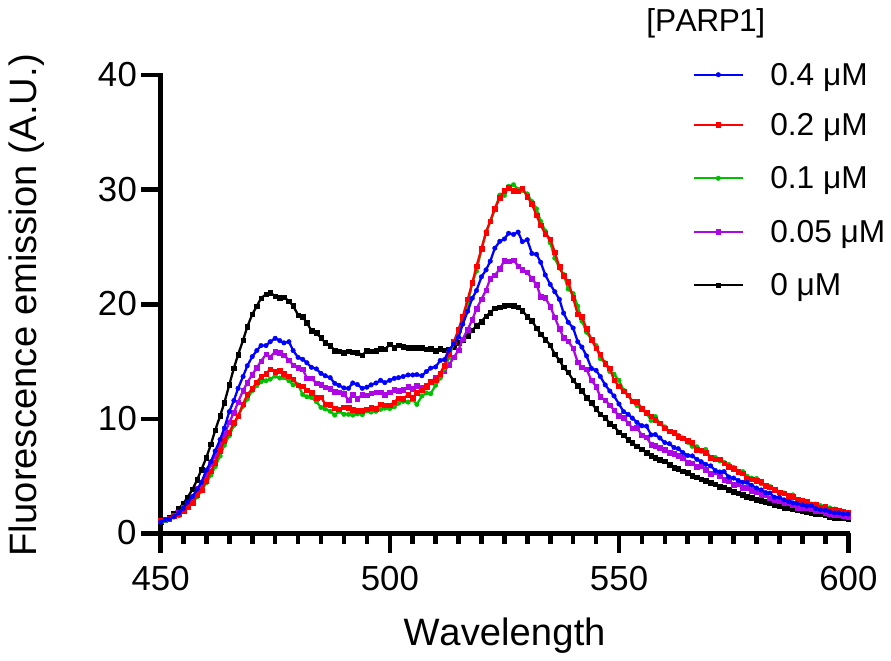

**Figure S1.** Fluorescence emission spectrum of pARS (5 μM) excited with 440 nm (+/- 10) light with different concentration of auto-modified PARP1. A ratio of 50 to 1 (pARS-to-PARP1) gives the best dynamic range.

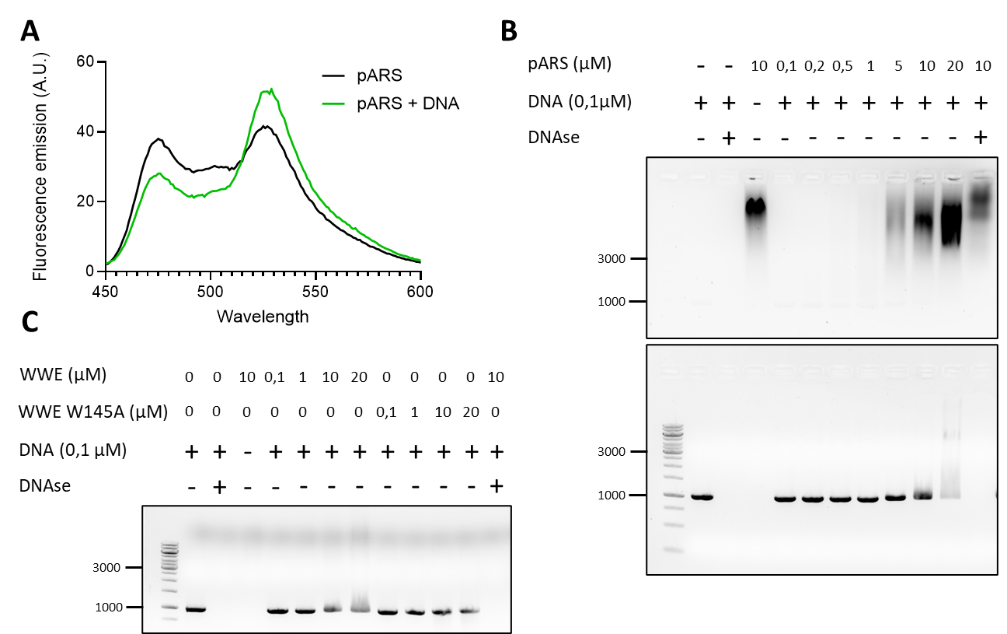

**Figure S2. A.** Fluorescence emission spectrum of pARS (5 μM) excited with 440 nm (+/- 10) light with or without DNA (0,1 mg/mL). **B.C.** DNA binding of pARS and WWE PBD were assessed with a dose response of respective purified proteins with 0,1 μM DNA (1 kb).

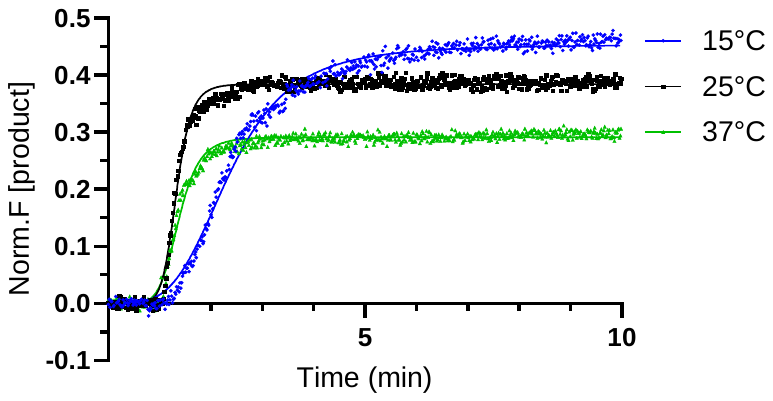

**Figure S3.** PARP1 (5 nM) dependent PAR formation under 10 μM NAD^+^ at 15, 25 or 37°C. Data is shown as mean ± SEM of two independent experiments.

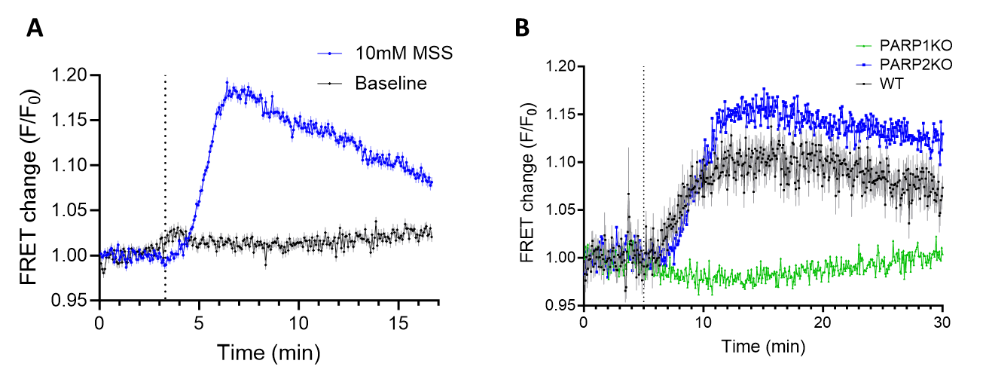

**Figure S4. A.** Average of 176 and 136 cell traces showing FRET change of the pARS sensor in HeLa cells after treatment with 10 mM MMS. Data is shown as mean ± SEM of three independent experiments. **B.** Average of 35, 55 and 43 cell traces showing FRET changes of the pARS sensor in U2OS cells WT, PARP1KO or PARP2KO after treatment with 10 mM MMS. Data is shown as mean ± SEM of two independent experiments.

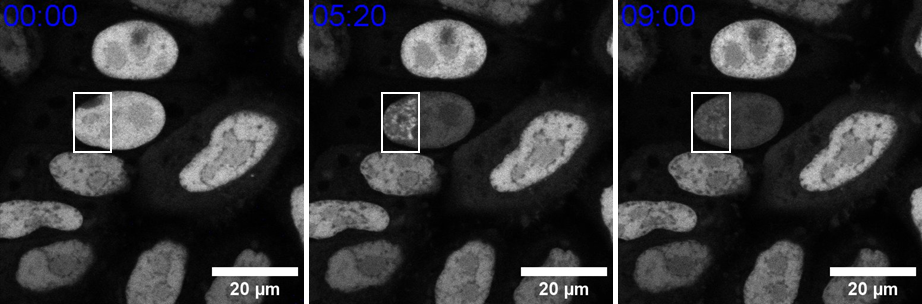

**Figure S5.** Confocal micrograph representing different time points of FRAP experiment using 375 nm laser at 100% laser power on HeLa Kyoto cells transfected with pARS (See supplemental video).

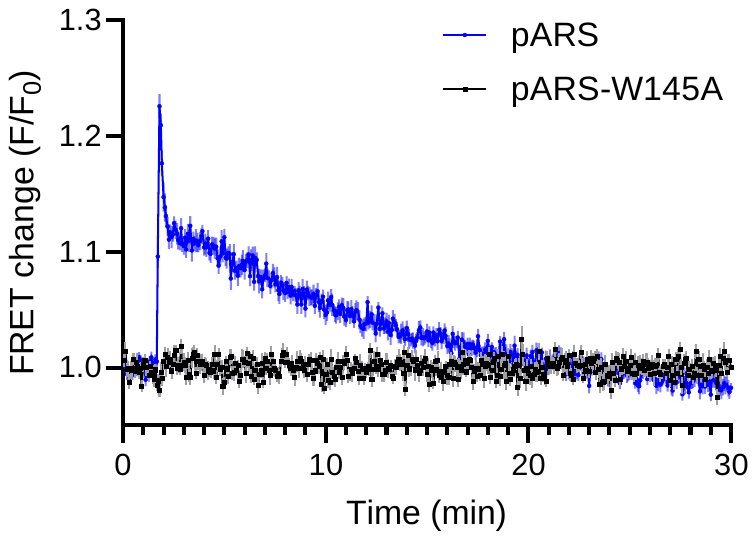

**Figure S6.** Average of 19 and 20 cell traces showing FRET changes of the pARS sensor or pARS-W145A in HEK 293 cells after irradiation with a 375 nm laser for 10sec. Data is shown as mean ± SEM of two independent experiments.

**MATERIALS AND METHOD**

**Chemicals**

Protease inhibitor cocktail (Roche Diagnostics), Mass Spectrometry grade proteases (Thermo Fisher), ADPr (Sigma), isoADPr (synthesize in this study), DB008 (provided by the Cohen lab)^[1]^, Olaparib (Selleck Chemical), PDD00017273 (SigmaAldrich), AZD5305 (Selleck Chemical), DNAse I (Roche Diagnostics), NAD^+^ (Sigma), Methyl methanesulfonate (Thermo Fisher), Hoechst (Thermo Fisher), activated DNA (Sigma).

**pARS overexpression and purification for *in-vitro* characterization**

pARS was overexpressed in *Escherichia coli* Rosetta (DE3) cells. A one-liter culture was grown at 37°C in LB-Miller broth containing kanamycin (50 μg/ml) with vigorous shaking (120 rpm) to an optical *A*600 of ∼0.7 before expression was induced by addition of IPTG to a final concentration of 0.5 mM. After further growth for 3 h, cells were harvested by centrifugation at 6000 × *g* for 20 min, and lysed by sonification in 40 ml of Buffer A [50 mM Tris (pH 7.5), 150 mM NaCl, 10mM beta mercaptoethanol, 1 U of DNase I and a tablet of cOmplete™ ULTRA protease inhibitor cocktail (Roche Diagnostics, Indianapolis, IL, USA)]. The cell lysate was cleared by centrifugation at 20 000 × *g* for 30 min at 4°C, and the supernatant was filtered using a 0.22 μm filter and loaded onto a 5 ml Ni-NTA column (Thermo, Hispur Ni-NTA) pre-equilibrated in Buffer A. Resin was then washed with 10 column volumes of Buffer A, 10 column volumes of Buffer B [50 mM Tris (pH 7.5), 10 mM imidazole, 150 mM NaCl, 10mM beta mercaptoethanol], followed by 100 ml of Buffer C [50 mM Tris (pH 7.5), 20 mM imidazole, 150 mM NaCl, 10mM beta mercaptoethanol] before the protein was eluted with Buffer C supplemented with 400 mM imidazole. The Ni-NTA eluate was reconcentrated using Amicon spin filter, filtered using 0.22 μm filter and further purified using a 5mL HiTrap Heparine HP column with a FPLC chromatography system (AKTA prime). The column was equilibrated with Buffer A and then sample was applied to the column with a flow rate of 3mL/min. Elution was done following the 3 steps program: 5 column volumes of Buffer A, 40 column volume of gradient from 0 to 100% of Buffer D [50 mM Tris (pH 7.5), 1 M NaCl, 10mM beta mercaptoethanol], and 5 column volume of Buffer D at a flow rate of 3mL/min. Fractions were analyzed on SDS-Page. Selected fraction was reconcentrated using Amicon spin filter using manufacturer protocol.

**Full spectrum acquisition and kinetics**

Excitation–emission spectra and kinetics were measured using a Cary Eclipse fluorescence Spectrophotometer (Agilent). Data acquisition was carried out in Hepes buffer (50 mM HEPES, 150 mM NaCl, 4 mM MgCl_2_ pH 7.5) supplemented with fresh 0.2 μM tris(2-carboxyethyl)phosphine (TCEP). For kinetics experiments PARP1 was added at 5 nM and pARS at 250 nM. The reaction was started after collection of a one-minute baseline by manual addition of NAD^+^ at various concentrations. Kinetics measurements were collected at one second intervals with excitation at 440 nm and emission at 475 and 527 nm under constant agitation. FRET change was calculated by dividing the acceptor fluorescence intensity by the donor fluorescence intensity and normalization to baseline. Maximum reaction rates were independently measured from the sigmoidal fit for each curve at different substrate concentrations. *Km* values were estimated using nonlinear regression analysis and curve fitting using the Michaelis–Menten function in Prism (GraphPad).

**In-vitro steady state PARP1-autoPARylation measurements with pARS**

PARP1 was added to a final concentration of 10 nM in Hepes buffer (50 mM HEPES, 150 mM NaCl, 4 mM MgCl_2_ pH 7.5) supplemented with fresh 0.2 μM TCEP. Inhibitors of various concentrations were pre-incubated before addition of NAD^+^ at 100 μM. The reaction proceeded at 30°C for 30 min before quenching of the reaction with 10 μM Olaparib. PARG (10 nM) or NUDT16 (3 μM) were added to the reaction mixture and incubated at 30°C for 90 min. pARS (0.5 μM) was added to the mixture and transferred to a 384 well plate for imaging on a Tecan plate reader using 440 excitation and sequentially collecting emission at 475 and 527 nm (+/- 10 nm). FRET change was calculated by dividing the acceptor fluorescence by the donor fluorescence and normalization to a condition without PARP1. IC_50_ values were estimated using three-parameter regression analysis and curve fitting with no further constraints using Prism.

**Binding assay agarose gel**

Recombinant PARP1 or WWE domain were incubated at various concentration with 1kb DNA (0.1 μM) at room temperature in 50 mM HEPES, 150 mM NaCl, 4 mM MgCl_2_ pH 7.5 for 5 min. The mixture was then loaded on a 1% Tris-acetate (fisher) agarose gel supplemented with ethidium bromide and ran for 20 min at 100 Volts. The gel was imaged on a BioRad imager in the UV channel (for DNA) and green fluorescence channel (for pARS).

**Stable cell line generation expressing pARS**

Flp-In-293 cells were purchased from Thermo Fisher Scientific (cat # R75007), cultured in DMEM supplemented by 10% FBS and grown at 37°C with 5% CO_2_. Stable expression of pARS was achieved by transient transfection of pARS and pOG44 using Lipofectamine 2000 (ThermoFisher Scientific, Cat # 11668019). To ensure sufficient genetic diversity, a 40% confluent cells in a T75 flask were transfected to ensure more than 10 colonies grew upon addition and selection with 100 µg/mL hygromycin (ThermoFisher Scientific, Cat #10687010). Following transfection and selection, all hygromycin-resistant HEK293 Flp-In cells were sorted for expression, expanded, and frozen for further experimentation. HEK293 Flp-In cells expressing pARS were maintained in 50 µg/mL hygromycin.

**Live cell imaging acquisition**

Cells were seeded in eight-well Lab-Tek microscope dishes for 24 h (to reach 60-70 % confluence) before transfection in full growth DMEM medium. 150 ng of the pARS sensor were mixed with 0.3 µL of JetPrime in 20 µL of the manufacturer buffer, incubated for 15 min and added to the cells. After overnight incubation the transfection medium was replaced with fresh full growth medium. 48 h later cells were imaged at 37°C in full medium. Imaging was performed on a dual scanner confocal microscope Olympus Fluoview 1200, using a 63x (oil) objective. The FRET sensor was excited using a 440 nm laser (at a laser power of 1.0%) and the signal was collected in the CFP/YFP emission channels. For DNA damage stimulation, cells were treated with 1 mM H_2_O_2_ or 10mM MMS or UV light irradiation. For the FRAP assay, cells were irradiated with 100% 375 nm laser for 4 sec. For full field of view irradiation (Figure5), cells were pre incubated with various concentration of Hoechst for 5 min then washed with fresh full growth medium before starting imaging. Cells were irradiated with ~20mW/cm2 for 1 second with the 375 nm laser.

**Microscopy image analysis**

All images were analyzed with FIJI software. Primarily, multi-channel images were separated into single channels and converted to 32-bit. Each channel was then smooth and the time course experiment was duplicated and stacked using the Z project function (using CFP channel). Using the stacked channel image, region of interest in the nucleus of each cell was manually defined. A ratio-metric image of the Venus channel divided by the CFP channel was then generated and the previously acquired ROI mask was superimposed to the time course experiment and the multi-measure function was applied to it. From this stack, we extracted mean single cell values from the time course experiment. Those values were then exported to an excel files for further analysis. The ratio was normalized to baseline and plotted on Graphpad.

**Statistics**

For experiment in cells, we report means ± standard errors. All the experiments were performed in biological triplicates except unless otherwise indicated. For *in-vitro* experiments, we report means ± standard errors unless specified otherwise in the figure legend.

**General Chemistry Experimental**

1H NMR were recorded on a Bruker DPX spectrometer at 400 MHz. Chemical shifts are reported as parts per million (ppm) downfield from an internal tetramethylsilane standard or solvent references as s (singlet), d (doublet), t (triplet), q (quartet), p (pentet), h (hextet), hep (heptet), m (multiplet), and br (broad) All reactions were run in flame or oven dried glassware under an atmosphere of dry argon unless otherwise noted Solvents were of ACS chemical grade (Fisher Scientific) and used without further purification unless otherwise indicated. Commercially available starting reagents were used without further purification. Analytical thin-layer chromatography was performed with silica gel 60 F254 glass plates (SiliCycle). Flash column chromatography was conducted with either pre-packed Redisep Rf normal/reverse phase columns Teledyne ISCO) or self-packed columns containing 200-400 mesh silica gel (SiliCycle) on a Combiflash Companion purification system (Teledyne ISCO). High performance liquid chromatography (HPLC) was performed on a Varian Prostar 210 (Agilent) with a flow rate of 20 ml/min using Polaris 5 C18-A columns (150 x 4.6 mm, 3 μm - analytical, 150 x 21.2 mm, 5 μm-preparative) (Agilent). HPLC analytical conditions: mobile phase (MP) A: 50 mM TEAB buffer (aq), mobile phase (MP) B: Acetonitrile; flow rate = 1.0 ml/min; UV-Vis detection: λ1 = 254 nm, λ2 = 220 nm. All final products were ≥95% purity as assessed by this method. Retention times (tR) and purity refer to UV detection at 220 nm. Low-resolution mass spectra were acquired on an Advion Mass-express.

**Synthesis of IsoADPr**

N^6^-benzoyl-9-[2-*O*-(2,3-di-*O*-acetyl-5-*O*-(di-*tert*-butyl)-phosphoryl-α-D-ribofuranosyl)-3-*O*-acetyl-5-*O*-(di-*tert*-butyl)-phosphoryl-β-D-ribofuranosyl]-adenine (2)

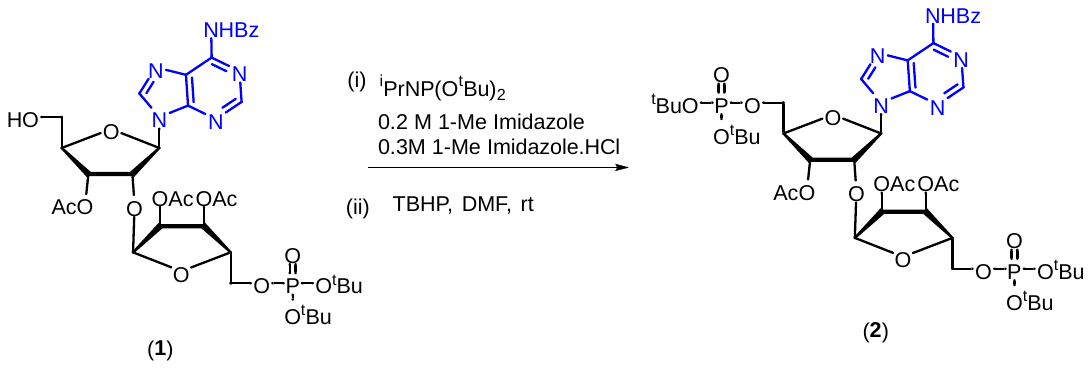

Compound 1 was prepared as reported in literature^[2]^. Compound 1 was co-evaporated with dioxane. To a 10 ml round bottom flask compound **1** (50.0 mg, 0.06 mmol) was taken and a mixture of 1-methyl imidazole.HCl (21.64 mg, 0.18 mmol) and 1-methyl Imidazole (9.99 mg, 0.12 mmol) was added. To this reaction mixture di-*tert*-butyl-*N*,*N*-di*iso*propylphosphoramidite (26.80 μL, 0.09 mmol) was added and the reaction mixture was stirred at room temperature for 30 min. Then mixtures was cooled to 0 °C and then TBHP in decane (60.84 μL, 5.5 M) was added. The solution was allowed to warm to room temperature and stirred for 1 h. The reaction was quenched by addition of aq. NaHCO_3_ and extracted with EtOAc. The organic layer was washed with water and dried over Na_2_SO_4_, concentrated *in-vacuo* and purified by flash silica gel chromatography using 0-5% DCM/MeOH as eluent to obtain the title compound **2** as a white foam (48.1 mg, 78%).

^1^H NMR (400 MHz, CDCl3) δ 9.06 (s, 1H), 8.81 (s, 1H), 8.37 (s, 1H), 8.05 – 7.98 (m, 2H), 7.64 – 7.58 (m, 1H), 7.53 (m, 2H), 6.27 (d, J = 6.0 Hz, 1H), 5.50 (dd, J = 5.2, 3.4 Hz, 1H), 5.30 – 5.28 (m, 1H), 5.26 (dd, J = 7.2, 3.2 Hz, 1H), 4.96 (t, J = 5.6 Hz, 1H), 4.76 (dd, J = 7.2, 4.6 Hz, 1H), 4.37 (p, J = 3.2 Hz, 1H), 4.24 (dd, J = 5.9, 3.2 Hz, 2H), 4.18 (p, J = 3.0 Hz, 1H), 4.04 (qdd, J = 11.3, 5.4, 3.1 Hz, 2H), 2.17 (s, 3H), 2.11 (s, 3H), 1.81 (s, 3H), 1.49 (d, J = 7.2 Hz, 18H), 1.44 (d, J = 4.0 Hz, 18H).

^13^C NMR (101 MHz, CDCl3) δ 170.0, 169.5, 169.40, 164.47, 152.9, 151.7, 149.6, 141.3, 133.7, 132.8, 128.9, 127.8, 123.0, 101.3, 86.2, 83.34, 83.30, 83.27, 83.23, 82.8, 82.7, 81.9, 81.8, 81.0, 80.9, 78.8, 71.9, 71.0, 69.7, 65.5, 65.49, 65.44, 29.9, 29.86, 29.81, 29.7, 20.8, 20.7, 20.1.

ESMS m/z calcd for C44H65N5O18P2 ([M+H]^+^) 1014.39, found 1014.11.

α-D-Ribofuranosyl-(1″→2′)-adenosine 5′,5″-bisphosphate triethylammonium salt (**3**, IsoADPr)

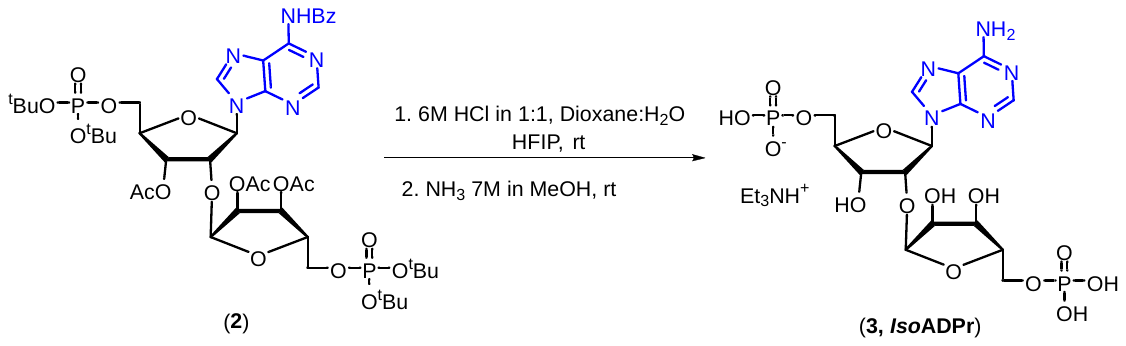

To a 10 ml round bottom flask containing **2** (20 mg, 19.72 µmol) was added 0.2 mL of HFIP (0.1M). After stirring for 1 min, a 6M solution of HCl (7.90 µL, 47.34 µmol) in 1:1 dioxane:water was added. The reaction was stirred at room temperature for 4 h. Solvent was evaporated *in-vacuo* and co-evaporated twice with NH_4_OH to obtain a crude residue. The latter was dissolved in ammonia solution (7M in MeOH, 0.5 mL) and the reaction was stirred for 24 h at room temperature. The solution was concentrated and triturated with acetonitrile. The white precipitate was purified by preparative HPLC (Gradient of 100% 50 mM TEAB buffer & 0% to 20% acetonitrile, 50 mM TEAB buffer & 80% acetonitrile). The fractions containing product were pooled and lyophilized to afford pure product **3** (6.1 mg, 56%). Spectral data were found to be in accordance as reported in the literature ^[3]^.

^1^H NMR (400 MHz, D2O) δ 8.54 (s, 1H), 8.27 (s, 1H), 6.30 (d, J = 5.5 Hz, 1H), 5.23 (s, 1H), 4.63 (s, 1H), 4.43 (s, 1H), 4.35 (s, 1H), 4.20 – 4.11 (m, 4H), 3.92 (s, 2H).

ESMS m/z calcd for C15H22N5O14P2 ([M-H]^-^) 558.06, found 558.06.

**^1^H NMR spectrum of compound 2**

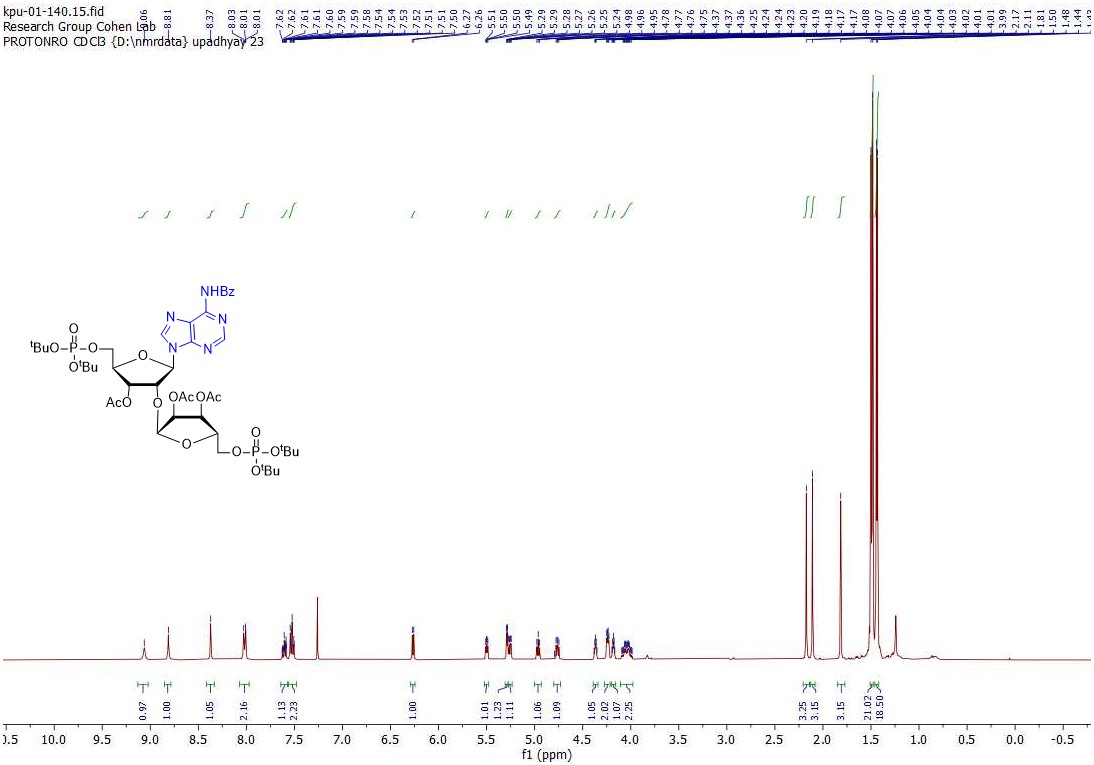

**13C NMR spectrum of compound 2**

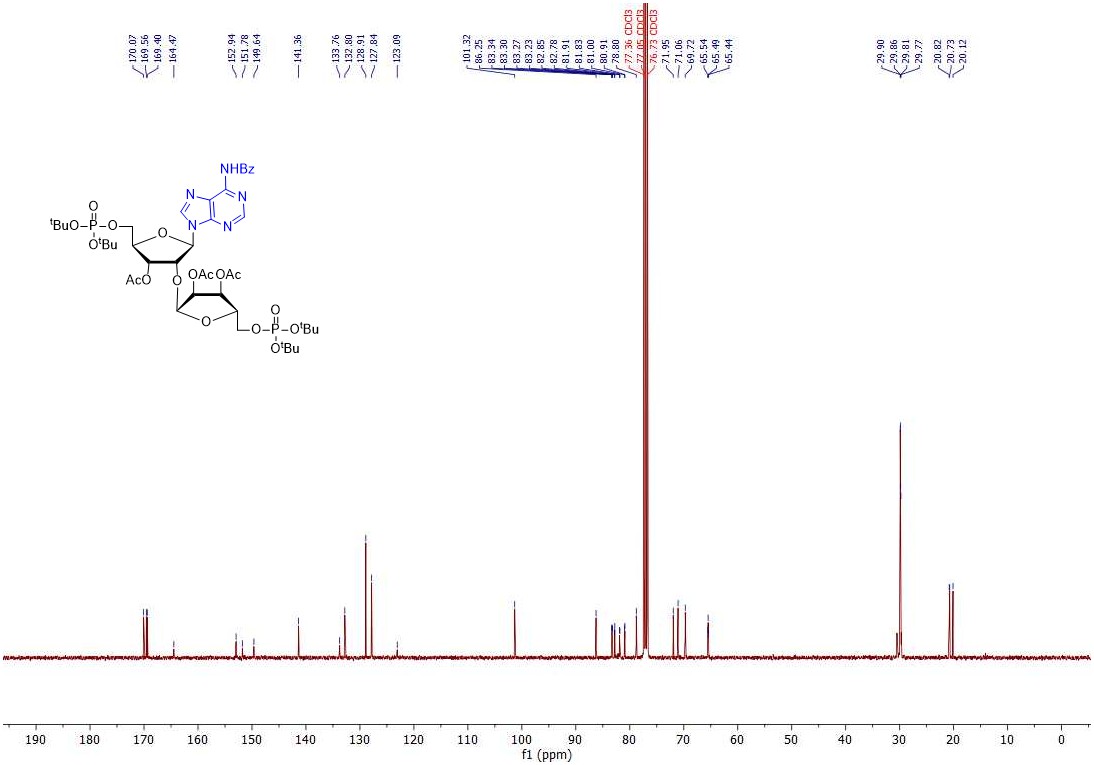

**1H NMR spectrum of compound 3, *Iso*ADPr**

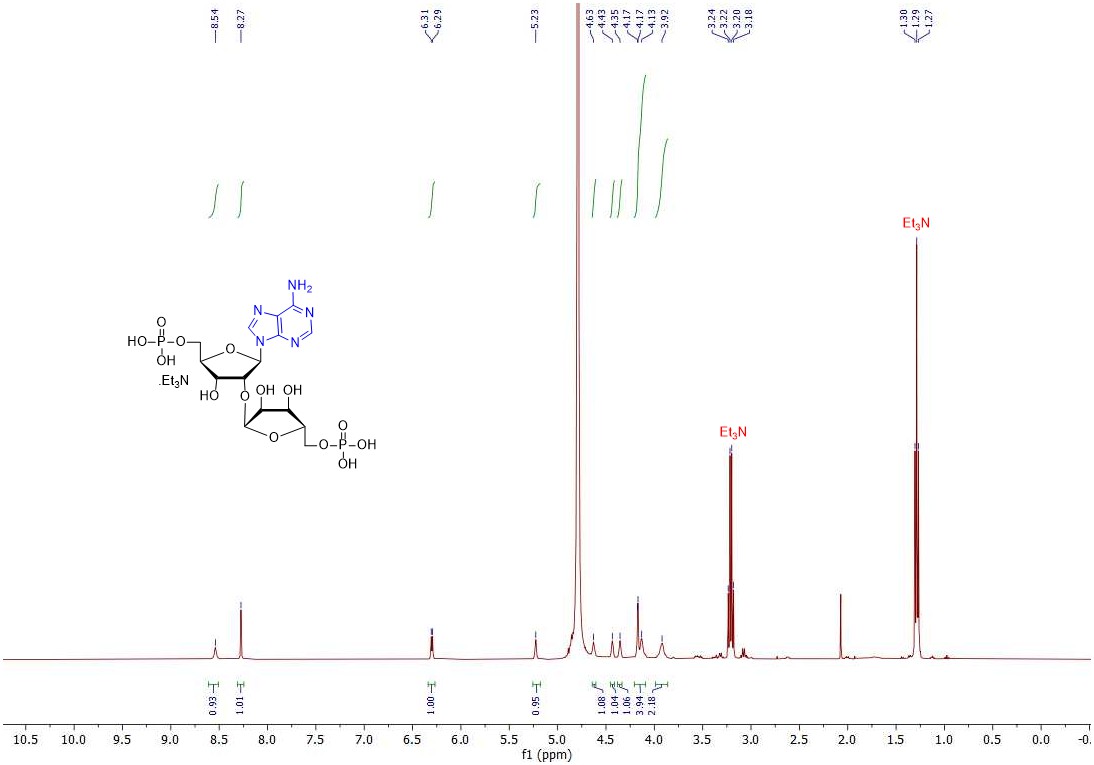
